# Structure and mechanism of potent bifunctional β-lactam- and homoserine lactone-degrading enzymes from marine microorganisms

**DOI:** 10.1101/2020.03.24.006742

**Authors:** Christopher Selleck, Marcelo Monteiro Pedroso, Liam Wilson, Stephan Krco, Esmée Gianna Knaven, Manfredi Miraula, Nataša Mitić, James A. Larrabee, Thomas Brück, Alice Clark, Luke W. Guddat, Gerhard Schenk

## Abstract

Genes that confer antibiotic resistance can rapidly be disseminated from one microorganism to another by mobile genetic elements, thus transferring resistance to previously susceptible bacterial strains. The misuse of antibiotics in health care and agriculture has provided a powerful evolutionary pressure to accelerate the spread of resistance genes, including those encoding β-lactamases. These are enzymes that are highly efficient in inactivating most of the commonly used β-lactam antibiotics. However, genes that confer antibiotic resistance are not only associated with pathogenic microorganisms, but are also found in non-pathogenic (*i.e*. environmental) microorganisms. Two recent examples are metal-dependent β-lactamases (MBLs) from the marine organisms *Novosphingobium pentaromativorans* and *Simiduia agarivorans*. Previous studies have demonstrated that their β-lactamase activity is comparable to those of well-known MBLs from pathogenic sources (*e.g*. NDM-1, AIM-1) but that they also possess efficient lactonase activity, an activity associated with quorum sensing. Here, we probed the structure and mechanism of these two enzymes using crystallographic, spectroscopic and fast kinetics techniques. Despite highly conserved active sites both enzymes demonstrate significant variations in their reaction mechanisms, highlighting both the extraordinary ability of MBLs to adapt to changing environmental conditions and the rather promiscuous acceptance of diverse substrates by these enzymes.

## Introduction

Antibiotic resistance is a rising socio-economic problem due to the lack of industrial antibiotic development over the last two decades^1,2^. Although new antibiotic substances are developed continuously by academic groups, very few are transferred into industrial production and clinical application^3^. Moreover, there is a lack of new (bio)chemical strategies to overcome antibiotic resistance, and although many of the newly developed antibiotic compounds may display novel structural features they mostly act upon the same cellular mechanisms (*i.e*. inhibition of cell wall biosynthesis, blocking ribosome function or cell replication). Therefore, there is an imminent global threat that available antibiotic substances may not be effective against ever faster evolving microbial pathogens; the most prominent among these pathogens are listed by the WHO and categorised in class 1-3 threat levels.^4^ To meet the societal challenges exerted by these microbial pathogens, new mechanisms to combat antibiotic resistance have to be identified, which can be the basis for the development of drugs that open new routes for anti-bacterial intervention.

More generally, antibiotic resistance has emerged in both Gram-positive and Gram-negative bacteria, and it is the latter group (in particular the ESKAPE pathogens, *i.e*. ***E****nterococcus faecium*, ***S****taphylococcus aureus*, ***K****lebsiella pneumoniae*, ***A****cinetobacter baumannii*, ***P****seudomonas aeruginosa* and the ***E****nterobacter* species) that poses an increased threat to public health, as reflected by the WHO prioritization. There are only a few antimicrobial agents available to battle these pathogens^1,2,5^, and it is not just a question of availability and diversity of these compounds, it is also a question of which cellular mechanisms they may target. In that respect, the current options are very limited and need urgent expansion^6,7^.

Sustained periods of haphazard use of antibiotics in both the medical sector and agriculture (*i.e*. the mass use of antibiotics in animal husbandry and plant growth), combined with enhanced global mobility, are pivotal factors driving the development of antibiotic resistance in both humans and domesticated farm animals^8^. The rapid spread of resistance is exemplified by a group of enzymes, metallo-β-lactamases (MBLs), which are highly efficient in deactivating nearly all types of naturally occuring β-lactam-based antibiotics (*e.g*. penicillin G, cephalosporins, carbapenems), and for which no clinically useful inhibitors are currently available^9-18^. However, although anthropogenic activities are a major contributor to the sharp rise in antibiotic resistance^19,20^, resistant microorganisms (many of which produce MBLs) have also been detected in pristine, unchallenged environments, such as the frozen tundra of a remote Alaskan region^19-22^. Therefore, studying MBL properties from “unchallenged”, *i.e*. pristine environments, may hold clues about their origin and evolution, which in turn can identify general microbial routes how these confer antibiotic resistance. Uncovering such routes or alternative roles of MBL in microbial interactions with their natural environment may, ultimately, provide strategies to develop new drugs that combat microbial infections that clearly constitute a major threat to modern health care.

MBLs belong to the family of binuclear metallohydrolases, a large group of enzymes that can accommodate two closely spaced metal ions in their active sites and which hydrolyze phosphoester and amide bonds in a broad range of substrates^9,10,14,15^. In the search for novel MBL-like proteins from non-pathogenic organisms, our group previously identified two candidates, one from *Novosphingobium pentaromativorans* and one from *Simiduia agarivorans*, both marine microorganisms^23,24^. The two proteins were initially labelled Maynooth Imipenemase 1 and 2 (*i.e*. MIM-1 and MIM-2^25^), a nomenclature associated with the place (Maynooth University, Ireland), where they were initially described and characterized. However, while both enzymes do belong to the B3 subgroup of the MBLs, they only share modest overall sequence similarity with each other (23%). They also display some distinct variations with respect to their preferred substrates^23,24^. Thus, in order to make sure that the two enzymes are not mistaken as variants of the same proteins we propose to maintain only the MIM-1 label but rename MIM-2 to *S. agarivorans* MBL-1, *i.e*. SAM-1. Both are efficient MBLs^24^, but are also very potent in hydrolyzing a range of homoserine lactone substrates^23^; the latter may suggest that these enzymes play an important role in quorum sensing. In order to further probe the function of MIM-1 and SAM-1, we have investigated their crystal structures and compared them with those of other MBLs. We also employed a range of spectroscopic and fast kinetics techniques to probe their reaction mechanism, and found evidence that further highlights the mechanistic flexibility, that is characteristic for at least some of the MBLs^26^.

## Materials and Methods

MIM-1 and SAM-1 were expressed and purified using a previously published procedure^23,24^. In brief, BL21(DE3) cells were transformed with the plasmid *bla*_*MIM-1*_ or *bla*_*SAM-1*_. The proteins were expressed in LB medium, supplemented with 50 µg/ml kanamycin. Initially, the cell cultures were grown at 37 °C until the OD_600_ reached 0.4 - 0.6. Expression was subsequently induced by the addition of 1 mM of IPTG at 18 °C. The cell culture was then grown for another 48 hours, after which the cells were harvested by centrifugation and purified on a Hi-trap Q FF column, equilibrated with 20 mM Hepes buffer, pH 7.5, 0.15 mM ZnCl_2_. Proteins were eluted with a linear gradient from 0 to 1 M NaCl. Fractions containing activity against cefuroxime were combined, concentrated and subsequently loaded onto a Hiprep 16-60 Sephacryl S-300 HR gel filtration column, and eluted with 50 mM Tris buffer, pH 7.2, containing 0.15 mM ZnCl_2_. The fractions were at least 97% pure, judged by SDS-PAGE gel analysis, and the purified protein was stored in 10% glycerol at −20 °C. The protein concentration was determined by measuring the absorption at 280 nm (ε = 36,815 M^-1^cm^−1^ and 41,285 M^−1^cm^−1^ per monomer for MIM-1 and SAM-1, respectively)^24^.

### Crystallization, X-ray diffraction data collection and refinement

Crystals were prepared using the hanging-drop diffusion method at 18 °C. The drop solution contained 300 µL of the desired enzyme (*i.e*. 40 mg/mL MIM-1 or SAM-1) and 300 µL of the precipitant buffer. For MIM-1 the precipitant buffer used was 0.05 M citrate, pH 5, 0.05 M BisTris propane, pH 9.7, and 16% w/v PEG-3350; for SAM-1 it was 0.1 mM DS56E8, 0.1 M sodium citrate, pH 5.5, and 22% w/v PEG-3350. Typically, diamond-shaped (MIM-1) or plate-like (SAM-1) crystals began to form after seven days and continued to grow for the next 6 days. Diffraction data were collected with cryo-protected crystals in a mixture of 20% glycerol added to the precipitant buffer.

Crystallographic data were collected by remote access on beamline MX-2 at the Australian Synchrotron (Melbourne) using BLU-ICE^27^. The data were integrated, scaled and merged using HKL-2000^28^. Refinement and model building were carried out using PHENIX 1.8.4^29^ and COOT 0.7^30^, respectively, using the previously published coordinates for the B3 MBL AIM-1 from *P. aeruginosa* (4P62)^31^. All atoms were subsequently refined with anisotropic B-factors; most hydrogen atoms were fitted as riding models, though the proton of the bridging hydroxide was added manually based on the electron density. Relevant crystallographic data and refinement statistics are summarized in **Table 1**.

**Table 1.** Crystallographic data and refinement statistics for MIM-1 and SAM-1.

| Diffraction data | MIM-1 | SAM-1 |
| --- | --- | --- |
| Resolution Range (Å) | 48.06- 2.6 | 42.83- 1.84 |
| Observations ($I > \sigma(I)$)<sup>a</sup> | 230230 (27609) | 216935 (9576) |
| Unique reflections ($I > \sigma(I)$) | 16484 (1964) | 24468 (1318) |
| Completeness (%) | 99.9 (99.6) | 99.3 (88.2) |
| Mean $\langle I/\sigma(I) \rangle$ | 21.8 (2.2) | 7.5 (2.2) |
| R<sub>merge</sub> | 0.0.078 (1.014) | 0.168 (0.536) |
| R<sub>pim</sub> | 0.022 (0.278) | 0.06 (0.205) |
| Multiplicity | 14.0 (14.1) | 8.9 (7.3) |
| Crystal parameters |  |  |
| Space group | P 41 21 2 | P 2 21 2 |
| Unit cell lengths (Å) | a = 67.96 b = 67.96 c = 216.67 | a = 51.67 b = 69.53 c = 76.54 |
| Unit cell angles (°) | $\alpha = 90 \beta = 90 \gamma = 90$ | $\alpha = 90 \beta = 90 \gamma = 90$ |
| Refinement |  |  |
| R<sub>work</sub><sup>b</sup> | 0.1728 | 0.2602 |
| R<sub>free</sub><sup>b</sup> | 0.2194 | 0.3090 |
| rmsd bond lengths (Å) | 0.0062 | 0.0019 |
| rmsd bond angles (°) | 0.860 | 0.568 |
| Ramachandran plot statistics |  |  |
| Favoured regions | 93.73 | 94.19 |
| Outlier regions | 0.37 | 1.55 |
| PDB code | 6AUF | 6MFI |

### Magnetic circular dichroism (MCD) spectroscopy

The preparation of the apo-enzyme was performed following a previously published procedure^26,32,33^ by adding 10 mM EDTA in 20 mM Hepes buffer, pH 7.0, to the purified enzyme preparations (concentrations for MIM-1 and SAM-1 were 12 mg/mL and 18 mg/mL, respectively). After 24 h incubation at 4 °C the chelating agent was removed using a desalting column (Econo-Pac 10DG from Bio-Rad), pre-equilibrated with 20 mM Hepes buffer, pH 7.0. Subsequently, three equivalents of Co^2+^ were added to the metal-free protein solution and the mixture was incubated over night at 4 °C. The excess of Co^2+^ was removed using a desalting column pre-equilibrated with 50 mM Tris.HCl buffer, pH 8.0.

For spectroscopic measurements this solution was diluted with glycerol to a final concentration of ∼1 mM (*i.e*. a 3:2 glycerol:buffer mixture). The samples were transferred to a 0.62 cm path length nickel-plated copper sample cell with quartz windows. The MCD system used has a JASCO J815 spectropolarimeter and an Oxford Instruments SM4000 cryostat/magnet. Data were initially collected at 7.0 T and 1.4 K. Variable-temperature, variable-field (VTVH) data were subsequently measured at increments of 0.5 T from 0 to 7.0 T and at temperatures of 1.4, 3, 6, 12, 24 and 48K. The experimental spectra were plotted as a function of wavenumbers and fitted to a minimum number of Gaussian peaks to achieve the final composite spectrum using the GRAMS AI software package^34^. The data were subsequently analyzed as described in detail elsewhere ^26,33,35-39^.

### Pre-steady state kinetics measurements

All pre-steady state kinetic measurements were carried out using an Applied Photophysics SX-18 spectrometer, coupled with a photodiode array detector. Data were collected under single turnover conditions, where the concentration of the enzyme (∼70 μM) was superior to that of the substrate nitrocefin (∼30 μM). The protein samples were prepared in 50 mM Tris-HCl, pH 8.5. Data were recorded at 25 °C over a period of 0.5 seconds. Data from at least eight reproducible experiments were collected, averaged and corrected for the instrument dead time (1.5 ms). The kinetic experiments were simulated using the program KINSIM using the mechanistic model presented in **Scheme 1**; experimental data were fitted using the program FITSIM^40-42^ (see text for more details).

**Scheme 1.**
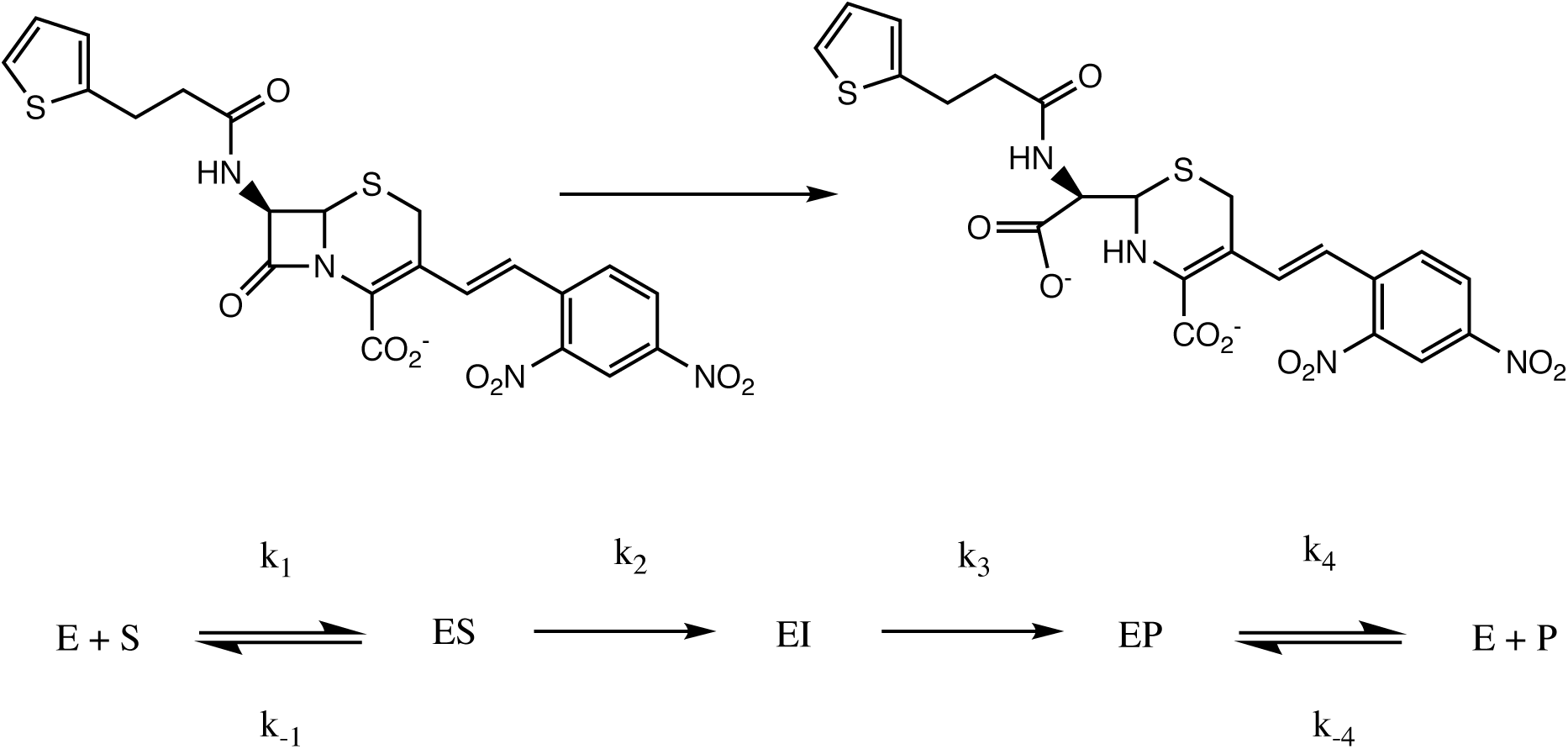
Hydrolysis of nitrocefin (top) and kinetic model (bottom) used for the modelling and fitting of the experimental data.

## Results and Discussion

### Protein purification and crystallography

Recombinant MIM-1 and SAM-1 were expressed in *E. coli* BL21(DE3) and purified using a previously protocol published ^23,24^. Importantly, the mature proteins had no N-terminal signal peptides, thus preventing their secretion into the periplasm. Both proteins crystallized over a period of approximately eight days.

For MIM-1, the large diamond-shaped crystals diffracted to 2.6 Å and belong to the space group P 41 21 2 (**Table 1**). Molecular replacement (using AIM-1 as a template) showed the presence of a single molecule occupying each asymmetric unit. The overall structure of MIM-1 consists of a well-defined electron density map containing 274 amino acid residues (out of 300 in total for the full-length enzyme), allowing for an uninterrupted trace of the polypeptide backbone from Pro33 to Ala305. Also present are the two catalytically important Zn^2+^ ions with occupancies of 1.0 each, along with a citrate molecule, which was a component of the crystallization buffer. The Ramachandran plot shows that most of the residues (93.73%) are within the favored regions, whilst another 1.97% are within the allowed region.

For SAM-1, plate-like crystals diffracted to a resolution of 1.8 Å and have the space group P 2 21 2, also with only one molecule per asymmetric unit (**Table 1**). The structure of SAM-1, solved using molecular replacement with AIM-1 as template, is well defined in the electron density map resolved for 264 residues (out of 295 for the full-length enzyme), allowing to trace of the polypeptide backbone from Thr31 to Ala300. However, the electron densities for the side chains of Ala160 to Leu161 and Glu207 to Arg210 are not clearly identifiable, and hence, these residues are not present in the final structure. Moreover two Zn^2+^ ions with an occupancy of 1.0 are present in the structure, which define the catalytically relevant active site. The Ramachandran plot indicates that 94.19% of all residues are within the favored regions whilst another 4.26% are within allowed regions. Further, the Patterson analysis function in Xtriage indicates a significant off-origin peak that is 20.8% of the magnitude of the origin peak. This observation strongly suggests the presence of pseudo-translational symmetry within the data. The refined overall structures of MIM-1 and SAM-1 are shown in **Figure 1**.

**Figure 1:**
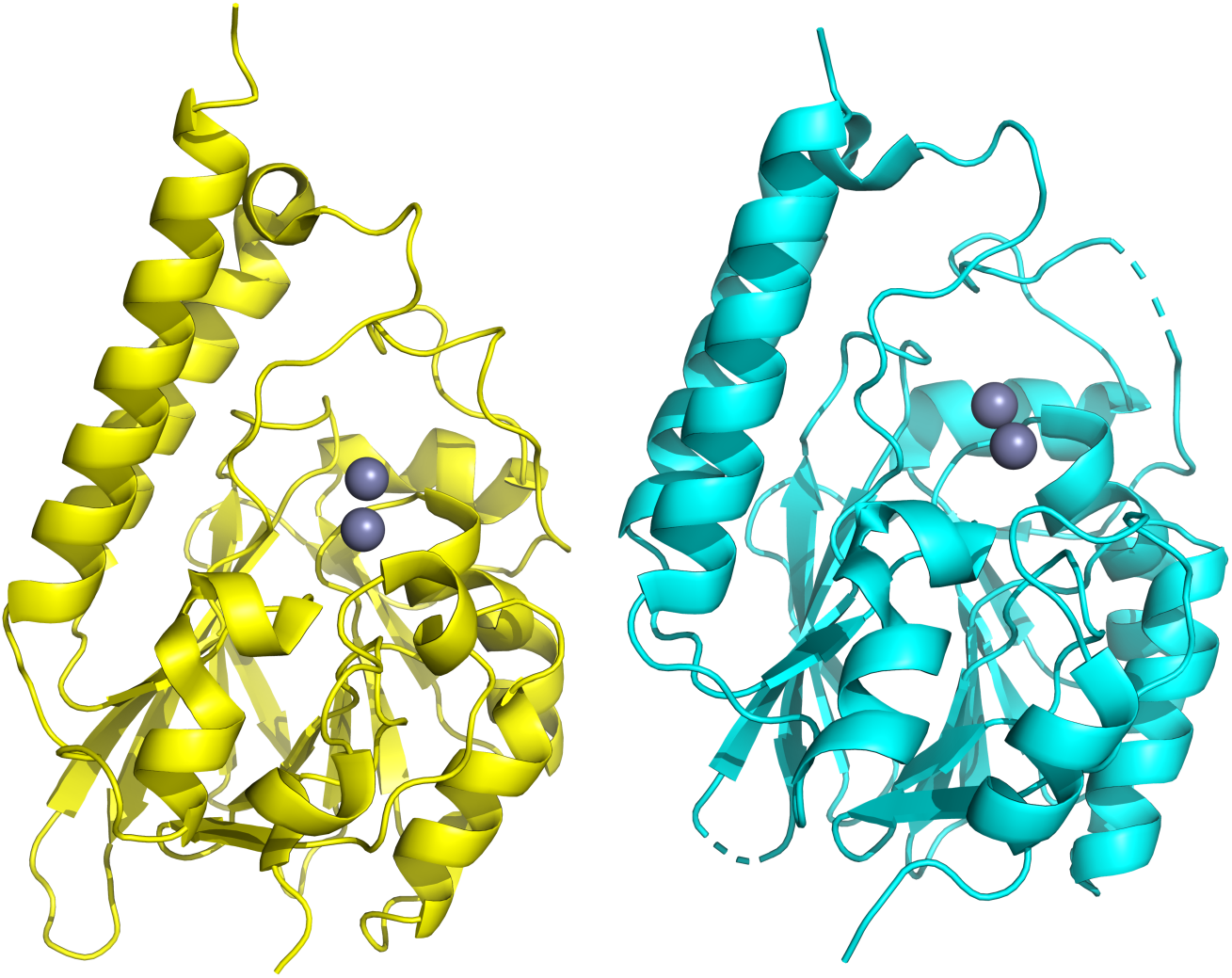
Overall structures of MIM-1 (yellow) and SAM-1 (cyan). Both enzymes contain the characteristic αβ/βα structural motif and are very similar to other MBLs from the B3 subgroup such as AIM-1^31^, L1^44^, SMB-1^45^ and BJP^46^. The catalytically relevant metal ion center is located in the middle. The Zn^2+^ ions are shown as grey spheres.

### Overall structure of MIM-1 and SAM-1

The overall folds of MIM-1 and SAM-1 consist of an αβ/βα motif, characteristic for MBLs, with the hydrophilic α helices exposed to the solvent and the central core formed by two β-sheets composed of five and seven β strands, respectively (**Figure 1**). Both MIM-1 and SAM-1 contain thirteen β-sheets with seven antiparallel strands at the N-terminus (B1-B7) and five antiparallel sheets at the C-terminus. Additionally, the MIM-1 structure also contains six α helices, two 3_10_ helices (η1 and η2) and a citrate molecule bound within the active site. In contrast, the SAM-1 structure contains 8 α helices and three 3_10_ helices (η1 to η3).

The active site groove in MBLs is defined by two loops, which are located at the interface of the two αβ domains and houses amino acid residues that are pivotal to the binding of both the catalytically essential Zn^2+^ ions and the substrates (**Figure 1**). Specifically, the two metal ions in the α and β sites (Zn1 and Zn2) of MIM-1 and SAM-1 are coordinated by His116, His118 and His194 (α site), and Asp120, His121 and His260/259 (β site), respectively. The motif HHH/DHH for the ligands interacting with the metals in the active site is a characteristic feature for MBLs belonging to the B3 subgroup^43^. Additional features in MIM-1 and SAM-1 include the presence of Gln157, located on loop 1 (**Figures 2** and **3**) and the presence of a disulphide bridge, which locks in the extended N-terminus, thus maintaining the active site pocket in an open conformation (**Figures 4** and **5**). The active site structures will be discussed in more detail in the following sections.

**Figure 2.**
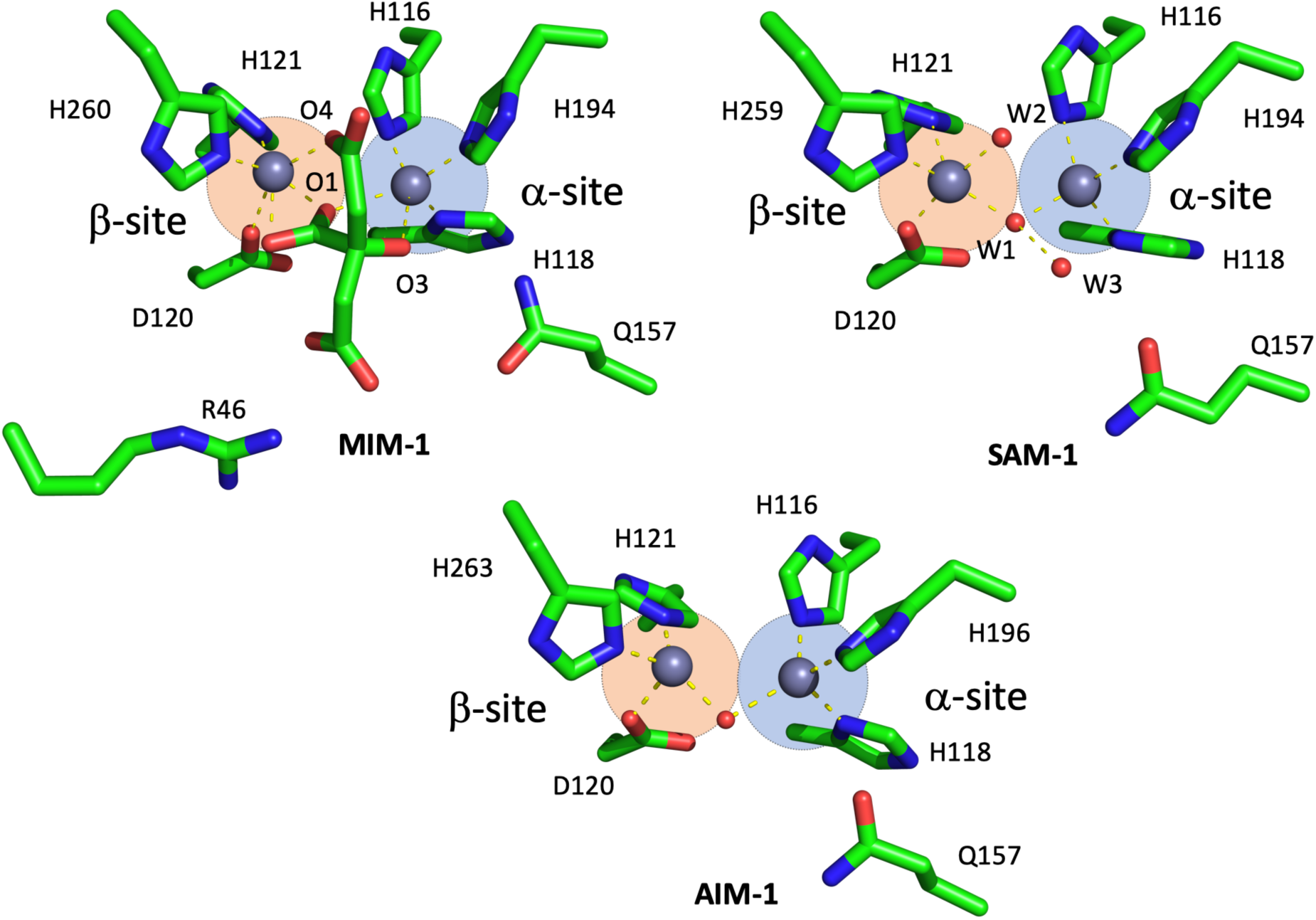
Active site structures of MIM-1 (left) and SAM-1 (right), including Gln157 in both enzymes and Arg46 (for MIM-1 only), which may play an important role in substrate binding. In MIM-1, a molecule of citrate (a component of the crystallization solution) is present in the active site; its oxygen atoms O1, O3 and O4 are replaced by water molecules (W1-W3) in the structure of SAM1.

**Figure 3.**
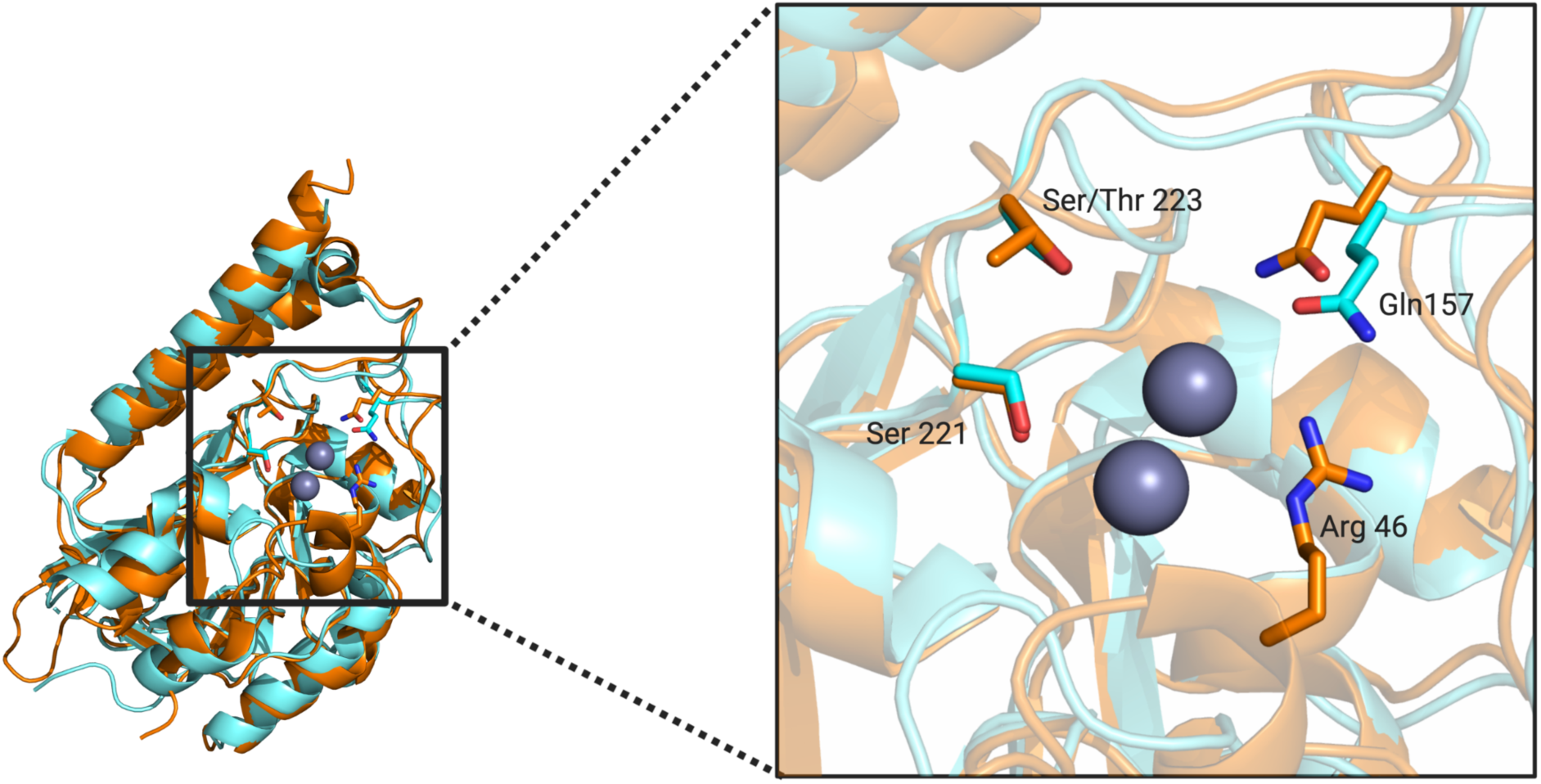
Loop 1 and 2 regions for MIM-1 (orange) and SAM-1 (blue). Important residues for substrate binding are also shown (see text for details).

**Figure 4.**
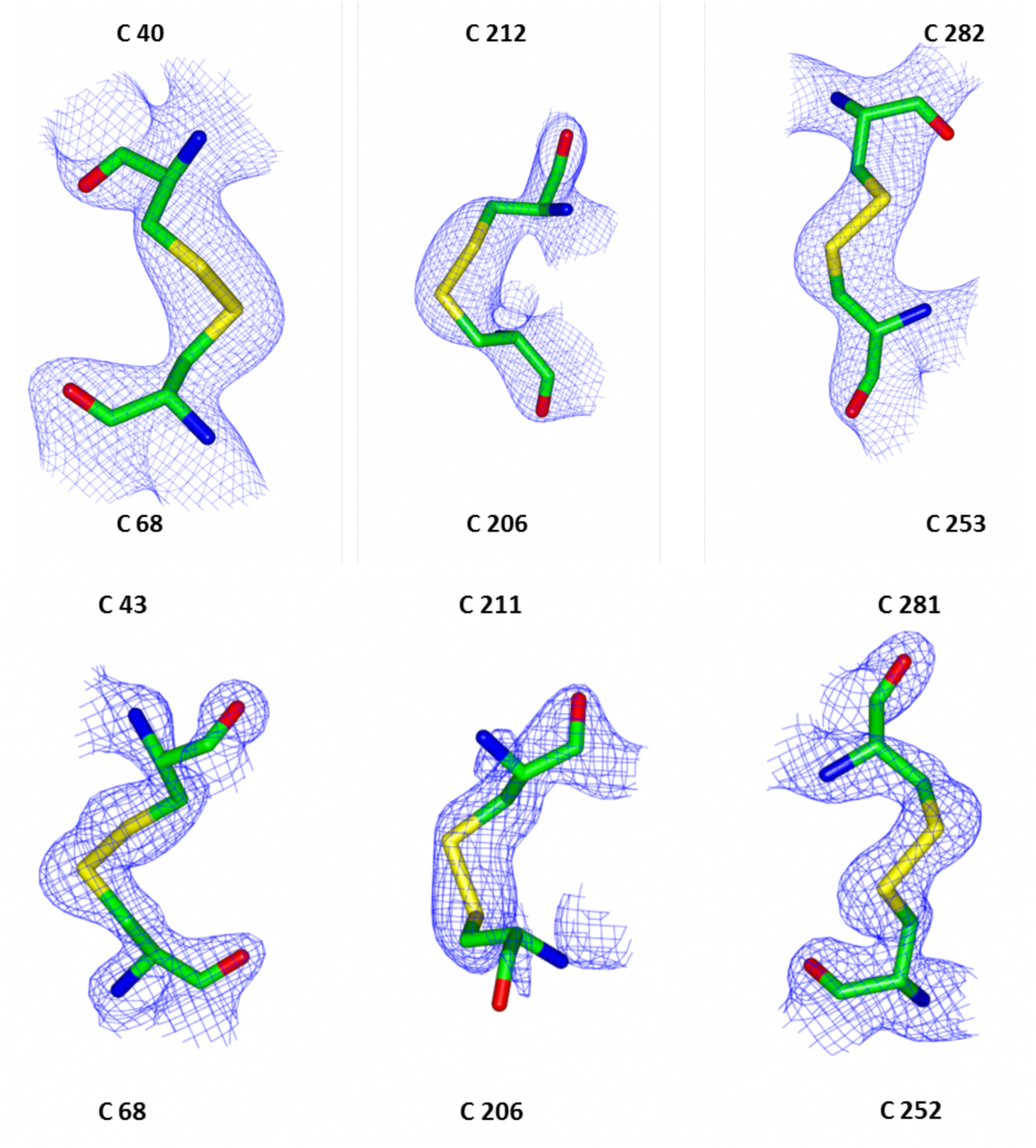
MIM-1 (top) and SAM-1 (bottom) have three disulfide bridges (shown with associated electron densities). Three disulfide bridges at equivalent positions are also present in AIM-1 and SMB-1, but fewer bridges are observed in other B3 MBLs such as BJP.

**Figure 5.**
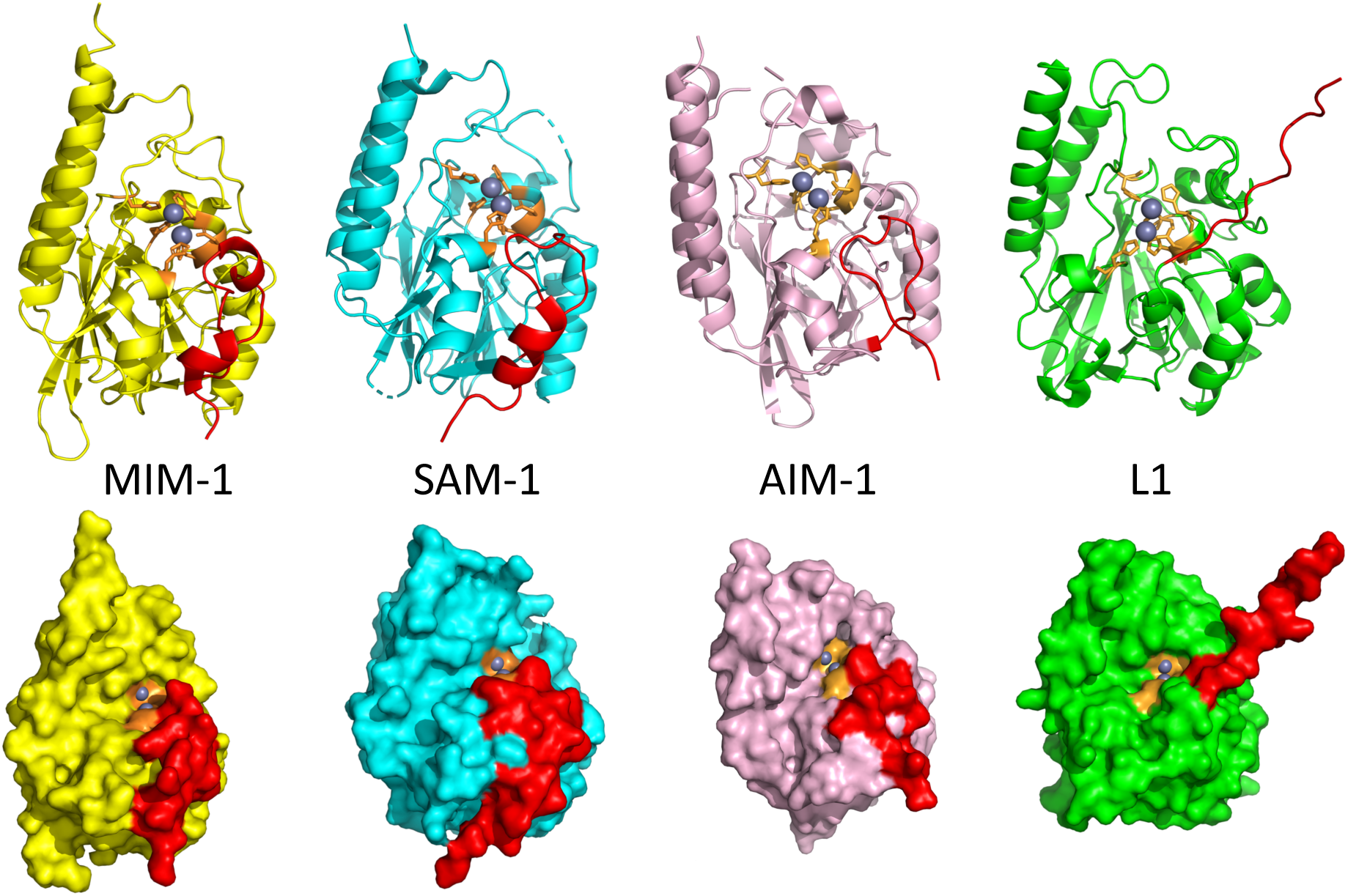
Selected crystal structures of MIM-1 (PDB 6AUF), SAM-1 (PDB 6MFI), AIM-1(PDB 4AWY) and L1 (PDB 1SML). The figure shows cartoon representations (top panel) and surface representation (bottom panel) of the overall structures of the selected enzymes, with active site ligands in orange, Zn^2+^ ions in grey, and the N-terminal loop in red. The extended N-terminus of L1 does not block the active site but may be involved in stabilizing the tetrameric form of the enzyme^44^. However, in MIM-1 and SAM-1 the Cys40-Cys68 and Cys43-Cys48 bridges, like in AIM-1, stabilize the extended N-terminus in a position that leaves access to the active site open.

### Zinc binding and active site

The zinc binding sites in MIM-1 and SAM-1 (**Figure 2**) are defined by a motif (HHH/DHH), that is characteristic for the majority of MBLs of the B3 subgroup^43-49^. The Zn-Zn distances in MIM-1 (4.02 Å) and SAM-1 (3.40 Å) are comparable with those observed in other B3 MBLs, *i.e*. BJP-1, FEZ-1, L1 and AIM-1^45-49^. The increased distance between the metal ions in MIM-1 is likely due to interactions with the bound citrate molecule, which is wedged within the active site.

Relevant distances in the metal binding sites of MIM-1 and SAM-1 are summarized in **Table 2**. Zn1 in the α site of SAM-1 adopts a distorted tetrahedral conformation with metal-ligand distances ranging from ∼2.0 to 2.2 Å. Apart from the three histidine ligands a water molecule (W1) completes the coordination sphere. A second water molecule (W2) is located in the vicinity. In MIM-1, this site resembles rather a distorted trigonal bipyramid with the three histidine ligands forming a triangular plane and two oxygen atoms of the bound citrate molecule (O3 and O1) occupying apical positions (**Figure 2**). In contrast, Zn2 in the β site adopts a distorted trigonal bipyramidal geometry in SAM-1 and an octahedral geometry in MIM-1. The plane of the bipyramid in the β site of SAM-1 is formed by ligands W1, W2 and His259, with ligands Asp120 and His121 occupying apical positions. Consequently, W1 in SAM-1 and oxygen atom O4 from bound citrate in MIM-1 are in metal-bridging positions. In the current models of the reaction mechanism employed by MBLs this position harbors the hydrolysis-initiating nucleophile^9,10,14^.

**Table 2.**
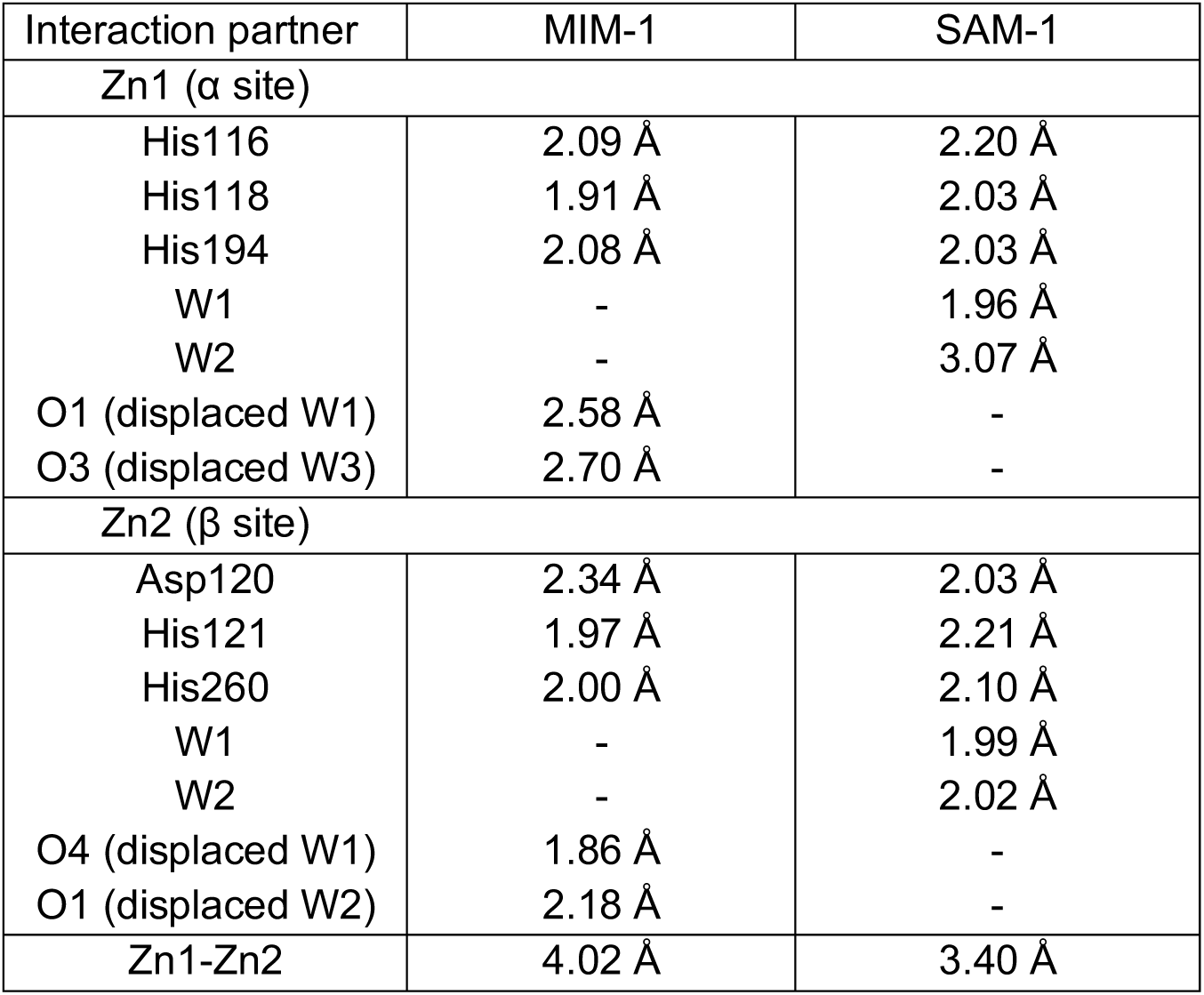
Selected distances between the two zinc ions in the active site of MIM-1 and SAM-1, and their ligands. W represents water molecules and O oxygen atoms of the citrate molecule bound to MIM-1.

| Interaction partner | MIM-1 | SAM-1 |
| --- | --- | --- |
| Zn1 ($\alpha$ site) | | |
| His116 | 2.09 Å | 2.20 Å |
| His118 | 1.91 Å | 2.03 Å |
| His194 | 2.08 Å | 2.03 Å |
| W1 | - | 1.96 Å |
| W2 | - | 3.07 Å |
| O1 (displaced W1) | 2.58 Å | - |
| O3 (displaced W3) | 2.70 Å | - |
| Zn2 ($\beta$ site) | | |
| Asp120 | 2.34 Å | 2.03 Å |
| His121 | 1.97 Å | 2.21 Å |
| His260 | 2.00 Å | 2.10 Å |
| W1 | - | 1.99 Å |
| W2 | - | 2.02 Å |
| O4 (displaced W1) | 1.86 Å | - |
| O1 (displaced W2) | 2.18 Å | - |
| Zn1-Zn2 | 4.02 Å | 3.40 Å |

In MIM-1, unique among MBLs, an arginine residue (Arg46), located on the hairpin loop leading toward the N-terminus, is in close proximity to the active site (**Figures 2** and **3**). Since this bulky and charged residue lines the channel that controls the access to the active center, it is possible that it may contribute to the recognition of substrates for this enzyme. However, despite some minor variations in the immediate vicinity of the catalytic centers the active site pockets of MBLs from the B3 subgroup are largely conserved, defined by two loops that house the residues responsible for substrate recognition and binding (**Figure 3**). Indeed, previous studies have illustrated that residues Gln157 (located on loop 1), Ser221 and Thr223 (both located on loop 2) play important roles in substrate recognition^48-50^. These residues are largely conserved in MIM-1, SAM-1, AIM-1 and SMB-1 (Thr223 is replaced by serine in MIM-1). Interestingly, it has been hypothesized that a glutamine in position 157 is only present in MBLs, that were acquired via horizontal gene transfer. In MBLs that are encoded on chromosomes (*i.e*. such as L1, BJP or FEZ-1) another residue is generally present in this position^44,45,47^. Thus, it is likely that both MIM-1 and SAM-1 are located on mobile genetic elements and were thus acquired through horizontal gene transfer.

The structural comparison between MIM-1, SAM-1 and AIM-1 indicates, that the three enzymes are very similar, especially in the vicinity of their catalytically relevant active sites. However, while each of these enzymes is a potent MBL only MIM-1 and SAM-1 are able to process homoserine lactones. We thus speculate, that the disulfide bridges present in the vicinity of the active sites of various B3 MBLs may play an important role in allowing homoserine lactone derivatives to bind in a catalytically competent conformation to the active site.

### MIM-1 and SAM-1 contain three disulfide bridges

Intramolecular disulfide bonds play an important role in protein folding and stability^48^. The majority of B3 MBLs possess at least one disulfide bond. In both MIM-1 and SAM-1 three disulfide bridges are present (**Figure 4**). The Cys253-Cys282 and Cys252-Cys281 pairs in MIM-1 and SAM-1, respectively, are present in the majority of characterized B3-type MBLs^48^. The two additional disulfide bonds (Cys40-Cys68, Cys206-Cys212 and Cys43-Cys68, Cys206-Cys211 in MIM-1 and SAM-1, respectively) are also present in AIM-1^48^ and SMB-1^49^ but no other B3-type MBLs. In particular, the Cys40-Cys68 (MIM-1) or Cys43-Cys48 (SAM-1) disulfide bridge directly affects the folding of the extended N-terminus in these enzymes. This bridge links the η1 helix with the β2 strand, causing the extended N-terminus to fold away from the active site (**Figure 5**). Other B3 MBLs, that share this structural feature (*i.e*. AIM-1 and SMB-1) tend to have higher MBL activity compared to those whose N-terminus enters or imposes on the space available within the active site (*i.e*. L1, BJP and Rm3)^44,45,49,51^. In agreement with this interpretation, it could be demonstrated, that the catalytic efficiency (*i.e*. the *k*_*cat*_/*K*_*m*_ ratio) of the B3 MBL L1 was enhanced at least 20-fold, when its extended N-terminus was truncated^52^. In MIM-1 and SAM-1 the catalytic efficiencies towards penicillin-based substrates and to some extent cephalosporins are comparable to those of AIM-1, but towards carbapenems MIM-1 and SAM-1 are one to two orders of magnitude less efficient than AIM-1^24^. In contrast, while MIM-1 and SAM-1 are efficient lactonases, AIM-1 is unable to hydrolyze such substrates^23^. Thus, while disulfide bridges undoubtedly aide the efficiency of some MBLs, they alone are not sufficient to determine the substrate preference of distinct members of this enzyme family. It appears, that MBLs involved in a pathogenic environment (*e.g*. AIM-1) have evolved to deal with the introduction of novel antibiotics, such as carbapenems. In contrast, related MBLs that are present in non-pathogenic organisms (*e.g*. MIM-1 or SAM-1) have adapted to accommodate substrates (*i.e*. quorum sensing mediators such as N-acyl homoserine lactones) that are used for different functions^23^. While our study has not yet led to the identification of specific residues or structural features, that validate the observed variations in substrate selectivity between MBLs such as AIM-1, MIM-1 and SAM-1 our study demonstrates that subtle factors may play a crucial role in determining the precise function of these potent enzymes.

### Spectroscopic characterization of MIM-1 and SAM-1 active site

Our crystallographic data demonstrate that MIM-1 and SAM-1, despite displaying some variation in substrate selectivity, are structurally very similar to other B3 MBLs, in particular to AIM-1. In order to investigate if there are any structural variations in solution, that may account for the observed functional differences we employed MCD spectroscopy. This method has been used extensively to probe the structure and mechanism of a series of metallohydrolases, including AIM-1^26,35-38,53-55^. Numerous studies, employed Co^2+^ derivatives of these enzymes to simplify data analysis^53^. In that context, Co^2+^ derivatives of MIM-1 and SAM-1 were generated previously and have been reported to be catalytically active^23,24^. Typical MCD spectra are shown in **Figure 6**; no fewer than five Gaussians are required to fit the data between 430-600 nm (**Table 3**).

**Table 3.**
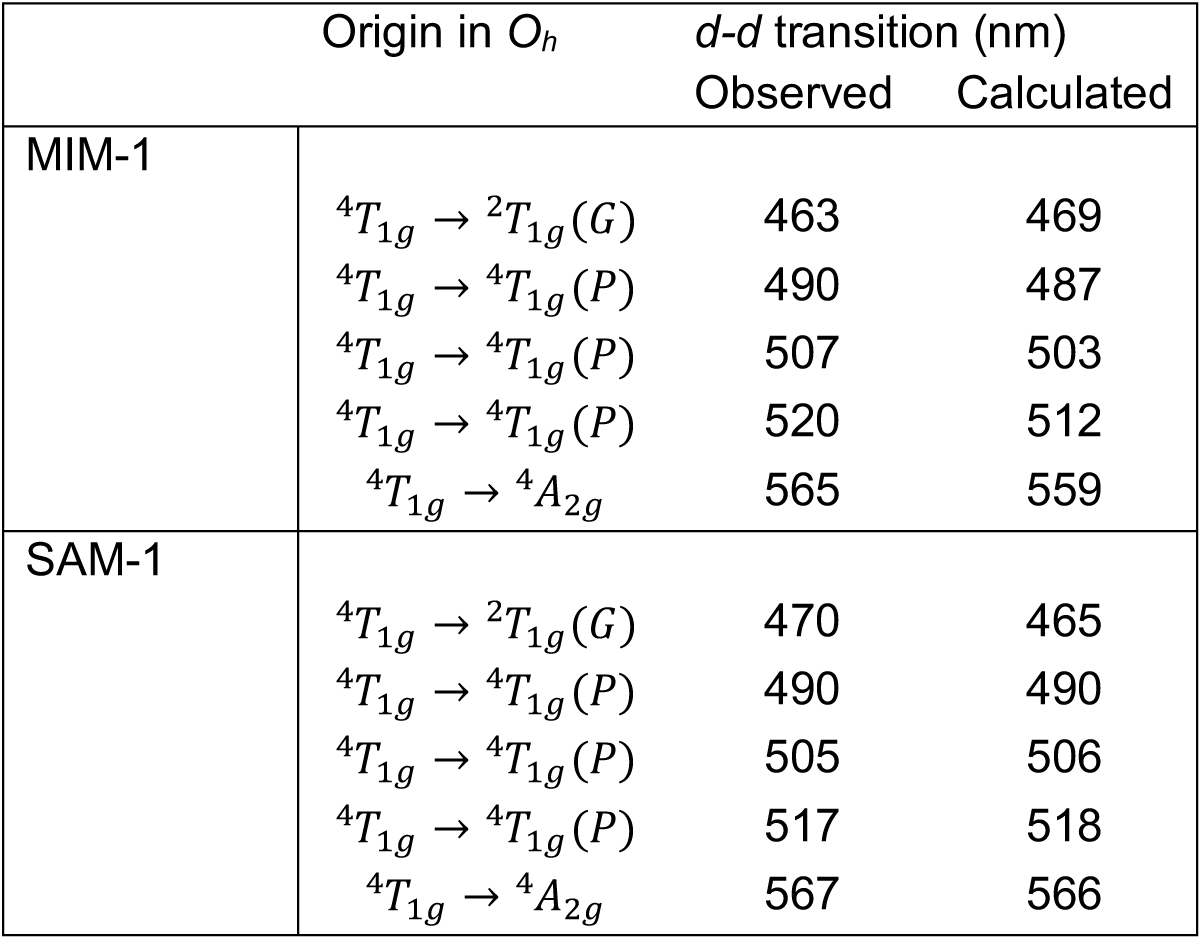
Summary of ligand field calculations

**Figure 6.**
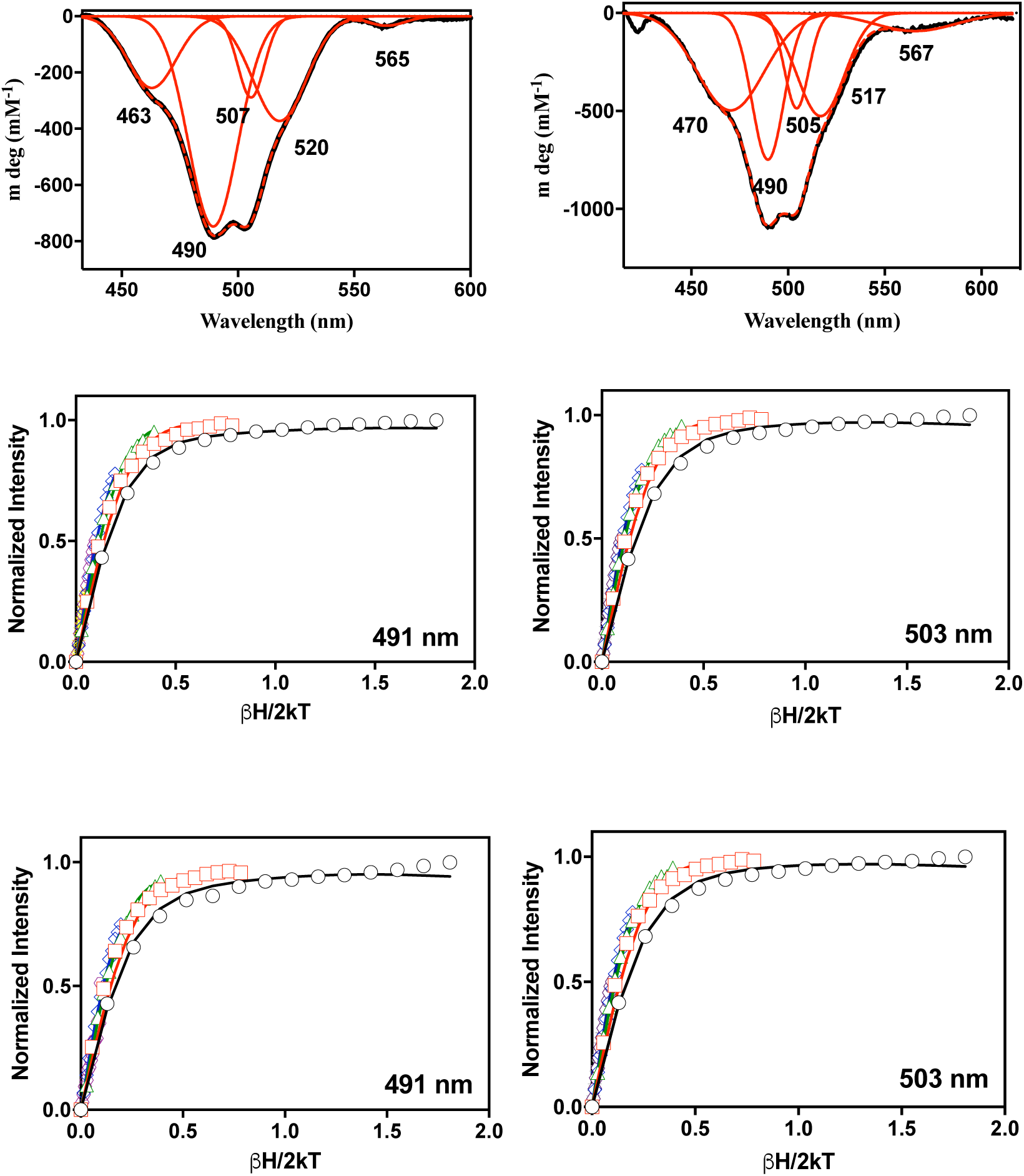
MCD spectra (at 1.4 K and 7 T) for MIM-1 (top left) and SAM-1 (top right) with Gaussian fits shown in red. The VTVH MCD data (middle panel for MIM-1 and bottom panel for SAM-1) for the 491 and 503/505 nm transitions. The isotherms shown in black, red, green, blue, yellow and grey represent data collected at 4.4, 3.0, 12.0, 24.0 and 48.0 K, respectively.

Calculations using the angular overlap model (AOM) were used to assign the individual transitions^53,56-58^. The coordinates applied for the calculation were obtained from the crystal structures of MIM-1 and SAM-1, and each Co^2+^ ion was treated independently. The Racah parameters *C* and *B* were fitted separately using the relation *C* = 4.7*B*. The analysis indicates that both Co^2+^ ions in the active sites of MIM-1 and SAM-1 are six-coordinate, contrasting with the crystallographic data obtained for the Zn^2+^ derivatives of these enzymes (**Figure 2** and **Table 2**). A similar observation was reported for AIM-1, for which it was suggested that water molecules complete the octahedral coordination environments of the two Co^2+^ centers^26^. Overall, the spectral data for AIM-1, MIM-1 and SAM-1 are similar, with AIM-1 having one extra transition around 530 nm. The electronic structures of the Co^2+^ ions were further analyzed by variable temperature, variable field (VTVH) MCD at three distinct transitions (*i.e*. 463 nm, 490 nm and 507 nm for MIM-1 and 470 nm, 490 nm and 505 nm for SAM-1; **Figure 6**). The data were analyzed using the dimer model as described elsewhere^21,26,33,35,38,53,54,59^ (**Table 4**). The bands at 490 and 507/505 nm are the main spin-allowed ^4^T_1g_ → ^4^T_1g_(P) transitions arising from the two six-coordinated Co^2+^ ions in the active site (as observed for AIM-1). Fitting parameters are summarized in **Table 4** and indicate that the two Co^2+^ have axial geometry (E/D ≈ 0) and that they are weakly ferromagnetically coupled *(J* ≈ 0.30 cm^−1^). While the metal ions in AIM-1 are also ferromagnetically coupled their exchange interaction is approximately three-fold weaker (*J* ≈ 0.10 cm^−1^)^26^. Thus, while the overall geometry in the active sites of AIM-1, MIM-1 and SAM-1 are similar, the significantly larger coupling interaction between the metal ions in the active sites of MIM-1 and SAM-1 may indicate some structural discrepancies that may align with the observed differences in substrate selectivity.

**Table 4.** Spectroscopic parameters obtained from fitting VTVH MCD data using the dimer model.

|  | MIM-1 |  |  | SAM-1 |  |  |
| --- | --- | --- | --- | --- | --- | --- |
| Parameter | 463 nm | 490 nm | 507 nm | 470 nm | 490 nm | 505 nm |
| J (cm<sup>-1</sup>) | 0.29 | 0.27 | 0.26 | 0.31 | 0.27 | 0.27 |
| D<sub>α</sub> (6-coordinate) (cm<sup>-1</sup>) | ≥ 50 | ≥ 50 | ≥ 50 | ≥ 50 | ≥ 50 | ≥ 50 |
| D<sub>β</sub> (6-coordinate) (cm<sup>-1</sup>) | - | - | - | - | - | - |
| E (α,β metal) (cm<sup>-1</sup>) | 0.0 | 0.0 | 0.0 | 0.0 | 0.0 | 0.0 |

### Rapid kinetics measurements

In a recent study, we employed stopped-flow absorbance measurements to probe the reaction mechanism of AIM-1^26^. Nitrocefin, a cephalosporin, was used as substrate since its hydrolysis can easily be observed due to the distinct spectroscopic properties of the intact molecule (λ_max_ = 390 nm; ε = 11,500 M^−1^ cm^−1^) and its hydrolyzed product (λ_max_ = 485 nm; ε = 17,420 M^−1^ cm^−1^). Furthermore, during the reaction a distinct ring-opened anionic intermediate (I) with a λ_max_ of 665 nm (ε = 32,000 M^−1^ cm^−1^) is formed^41,60^. In some MBLs such as NDM-1 and L1, the protonation of this reaction intermediate (*i.e*. the step EI→EP in **Scheme 1**) is rate-limiting, whereas in others (*e.g*. Bla2) the intermediate is too short-lived to be observed experimentally^25,60^. In AIM-1, the intermediate is observed but since its decay (∼22 s^−1^) is considerably slower than the recorded state-state catalytic rate for nitrocefin hydrolysis (240 s^−1^), it was concluded, that AIM-1 can employ two mechanistic strategies, depending on how the substrate is bound to the active site^26^.

Here, we employed similar experimental conditions as used for equivalent measurements with AIM-1. In a routine measurement, 30 μM of MIM-1 or SAM-1 were mixed with 70 μM of nitrocefin, and the progress of the reaction was monitored with a photodiode array detector between 320-800 nm (**Figure 7**). The profile for the reaction with MIM-1 resembles that recorded for AIM-1. The gradual depletion of the substrate is accompanied by a rapid emergence of the transient intermediate, which reaches a maximum concentration in under 50 ms. The intermediate then gradually disappears as the final product accumulates. The reaction traces of the substrate, intermediate and product were simulated with KINSIM^40,62,63^ using the model shown in **Scheme 1** to obtain estimates for the various rate constants. These values were then used to fit the data with the program FITSIM^40^. For the numeric evaluation, several simplifications were applied. Firstly, the formations of the enzyme-substrate (ES; *k*_*1*_) or enzyme-product (EP; *k*_*-4*_) complexes were assumed to be diffusion-limited and were thus locked in at 10^8^ M^−1^ s^−1^. Secondly, values for the constants *k*_*-2*_ and *k*_*-3*_ were set to zero since the hydrolysis of the β-lactam ring is essentially irreversible. Furthermore, initial values for the constants *k*_*2*_ and *k*_*3*_ were obtained from the rates of nitrocefin depletion and product formation, respectively. The simulations indicated, that the apparent reaction rate (k_app_) is not sensitive to the magnitude of *k*_*1*_, *k*_*4*_ and *k*_*-4*_, but strongly depends on the values of *k*_*-1*_, *k*_*2*_ and *k*_*3*_. The rate constants obtained though the appropriate data fitting are summarized in **Table 5**. In accordance to data obtained for AIM-1, the decay of the intermediate is the rate-determining step (*k*_*3*_) in the overall reaction of MIM-1. The microscopic rate constants summarized in **Table 5** can be used to calculate theoretical values for the steady-state parameters *k*_*cat*_ and *K*_*M*_^64^. For both MIM-1 and SAM-1 the calculated values of *k*_*cat*_ are similar to *k*_*3*_. However, only in MIM-1 a good agreement between the theoretical and experimentally determined *k*_*cat*_ values could be obtained (∼15 s^−1^ *vs* ∼17 s^−1^). Thus, MIM-1 is likely to employ the same mechanistic strategy as AIM-1, L1 and some biomimetic model complexes with the protonation of the anionic reaction intermediate being rate-limiting^9,26,63^.

**Table 5.** Rate constants for the reaction of MIM-1 and SAM-1 with nitrocefin. The rate constants were obtained using the mechanistic model illustrated in Scheme 1. *k*_*cat*_ and *K*_*M*_ values obtained from pre-steady (*i.e*. calculated using the King-Altman (KA) approach) and steady state (*i.e*. Michaelis-Menten (MM) kinetics) analyses are also included.

| Pre-steady-state | MIM-1 | SAM-1 |
| --- | --- | --- |
| $k_1 (\text{M}^{-1} \text{ s}^{-1})$ | $1.0 \times 10^8$ (fixed) | $1.0 \times 10^8$ (fixed) |
| $k_{-1} (\text{s}^{-1})$ | $1.0 \times 10^4$ | $2.0 \times 10^2$ |
| $k_2 (\text{s}^{-1})$ | $80 \pm 13$ | $5000 \pm 100$ |
| $k_3 (\text{s}^{-1})$ | $18 \pm 9$ | $100 \pm 2$ |
| $k_4 (\text{s}^{-1})$ | $2.3 \pm 0.1 \times 10^4$ | $1.7 \pm 0.1 \times 10^4$ |
| $k_{-4} (\text{M}^{-1} \text{ s}^{-1})$ | $1.0 \times 10^8$ (fixed) | $1.0 \times 10^8$ (fixed) |
| Macroscopic catalytic parameters (KA) |  |  |
| $k_{cat} (\text{s}^{-1})$ | $14.6 \pm 3$ | $97 \pm 6$ |
| $K_M (\mu\text{M})$ | $18 \pm 7$ | $25 \pm 0.4$ |
| Michaelis-Menten catalytic parameters (MM) |  |  |
| $k_{cat} (\text{s}^{-1})$ | $17 \pm 4$ | $10 \pm 5$ |
| $K_M (\mu\text{M})$ | $41 \pm 10$ | $20 \pm 3$ |

**Figure 7.**
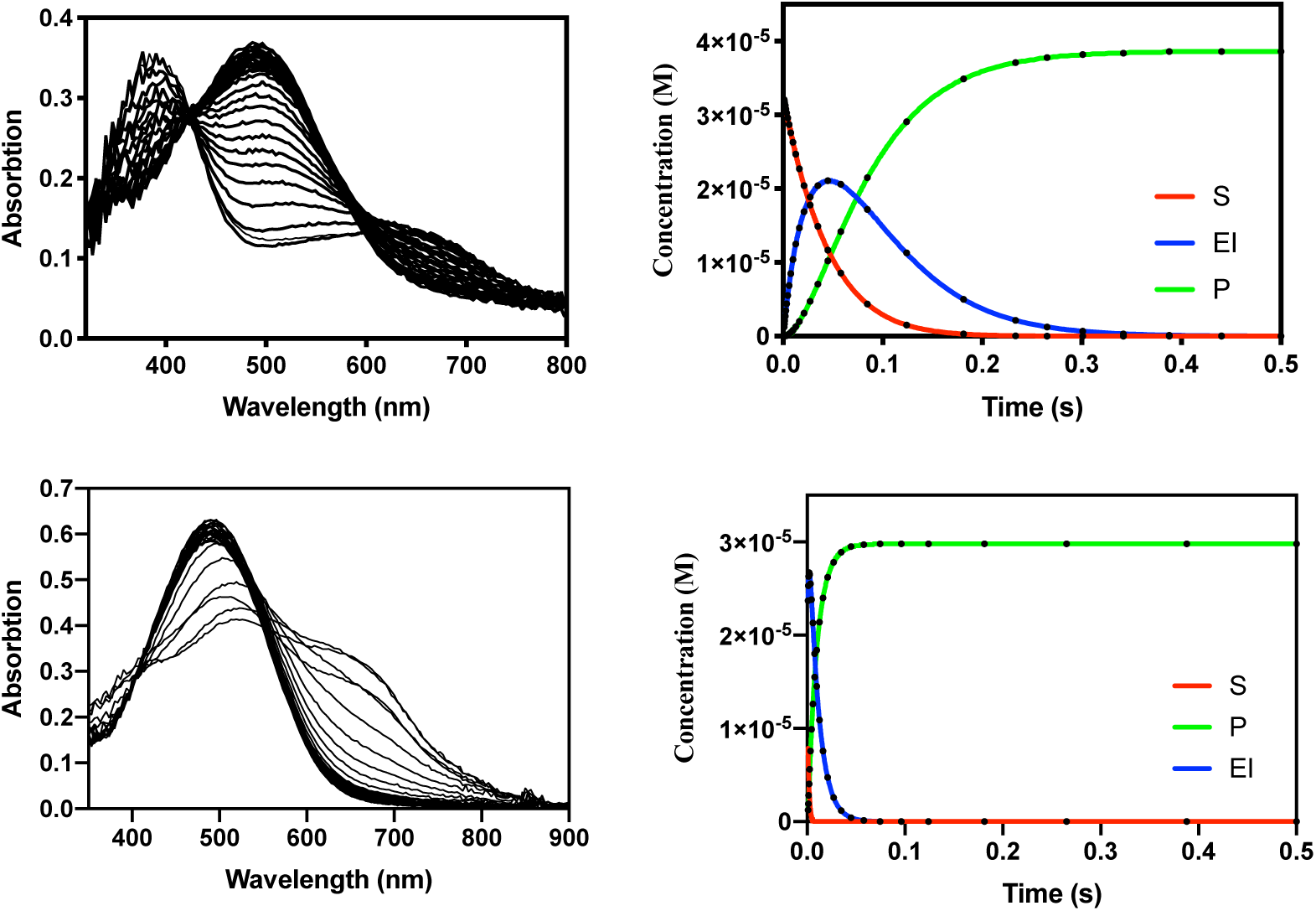
Stopped-flow data analysis for the experimental data for MIM-1 (top) and SAM-1 (bottom). The data were analyzed using the mechanistic model presented in Scheme 1 and the corresponding fits to the time course for the substrate, product and intermediate are shown in red, green and blue.

For SAM-1 the reaction profile differs significantly from that of MIM-1 or AIM-1 (**Figure 7**). Only two transients are observed, at 485 nm and 665 nm. Similarly, in *Bla2* also only two transients were observed albeit at 390 nm and 485 nm^26^. Thus, while in *Bla2* no anionic reaction intermediate may be formed during the reaction, in SAM-1 the substrate is consumed at a faster rate than the instrument dead time (∼1 ms), while the formed intermediate decays too rapidly to be monitored. Hence, only the intermediate decay is observed in the experiment. Consequently, the product of the reaction is also rapidly formed. The same simulation and fitting approach as for MIM-1 was used to analyze the corresponding data for SAM-1. As expected *k*_*2*_ (formation rate for the intermediate) is ∼60 times faster than in MIM-1 (**Table 5**). The value of *k*_*3*_ (*i.e*. rate of product formation) is 100 s^−1^, about five times faster than the corresponding rate in MIM-1 (18 s^−1^). Similarly, the macroscopic *k*_*cat*_ value (97 s^−1^) of SAM-1 is significantly higher than that of MIM-1 (∼15 s^−1^). However, while in MIM-1 a good agreement between the theoretical and steady-state catalytic parameters is observed, for SAM-1 similar magnitudes are only observed for the *K*_*M*_ values (25 µM *vs* 20 µM for the theoretical and experimental values, respectively). In contrast, the macroscopic *k*_*cat*_ value of SAM-1 (97 s^−1^) is nearly 10-times larger than the corresponding steady-state rate constant (10 s^−1^). Hence, while SAM-1 is able to hydrolyze the substrate much faster than MIM-1 the regeneration of the active and/or the release of the product appear to be rate-limiting in SAM-1. The fitting of the data is not sensitive to the magnitude of *k*_*4*_ and hence the two possibilities cannot be distinguished at present. However, considering the similar *K*_*M*_ values obtained for MIM-1 and SAM-1 in the reaction with nitrocefin it appears, that the interactions between these enzymes and the reactant are also similar. In consequence, we suggest that the regeneration of the hydrolysis-initiating nucleophile (*i.e*. water molecule W1 in **Fig. 2**) may be rate-limiting for SAM-1. This data interpretation is consistent with considerable the difference in the coordination geometry observed in the α site of these two enzymes.

In summary, we previously demonstrated that marine environments harbor a large number of MBLs from the B3 subgroup^43^. Of particular interest among those MBLs are those from the marine organisms *N. pentaromativorans* (MIM-1) and *S. agarivorans* (SAM-1, formerly known as MIM-2) as we previously already demonstrated that they are not only potent MBLs, but are also capable to hydrolyze quorum-sensing molecules^23,24^. Both MIM-1 and SAM-1 have a fold characteristic of MBLs and their active sites accommodate two closely spaced Zn^2+^ ions, surrounded by six invariant ligands. Notably, both enzymes have residue Gln157 in the vicinity of their active sites, a residue that only appears to present in MBLs that are located on mobile genetic elements, and which may thus indicate that MIM-1 and SAM-1 were acquired through horizontal gene transfer^48^. This raises the specter that marine environments may present a fertile reservoir for antibiotic-degrading activities. Enzymes, such as MIM-1 and SAM-1, while likely to play an important role in sensing in their natural environment, do have the capability to transmit an antibiotic resistance agent, that has the potential to be harmful to both public health and the food chain.

The design of effective inhibitors for MBLs has been a popular strategy to prolong the clinical use of long-term established compounds such as penicillin. While a large and growing number of leads have been developed none of them has yet reached a clinical stage^3,11,65-69^. It is thus of significant interest to find out why marine bacteria such as *N. pentaromativorans* and *S. agarivorans* harbor MBLs. A better understanding of the natural role(s) of MIM-1 and SAM-1 may facilitate the design of new preventative measures that can hinder pathogens to enter our human ecosystem, where they can cause disease. For instance, compounds that disrupt quorum sensing may hinder bacteria to acquire nutrients, thus preventing their propagation to form colonies that can initiate the formation of bacterial biofilms, which are a major effector in propagating infections, particularly in the clinical environment^70,71^.

## Acknowledgements

The authors would like to thank the National Health and Medical Research council of Australia (APP1084778) and Science Foundation Ireland (SFI/09/YI/B1756) for financial support. JAL also wishes to acknowledge the National Science Foundation (USA) for support from grants CHE-1904005, 1303852, and 0820965 (MCD instrument). TB gratefully acknowledges funding by the Werner Siemens foundation for establishing the new research field of Synthetic Biotechnology at the Technical University of Munich.

## Contributions

C.S, L.W., S.K., E.G.K., M.M. and M.M.P. were involved with data acquisition. M.M.P., N.M., J.A.L., L.W.G and G.S. were involved with the concept and design of the experiments. C.S., M.M.P., N.M., T.B., A.C., L.W.G and G.S. analyzed and interpreted the data. C.S., M.M.P., T.B., A.C., and G.S. drafted the manuscript. Critical revision of the manuscript was done by all authors.

